# Connecting Boolean Integrase Logic gates to a novel alkane control signal via engineered level matching

**DOI:** 10.1101/031542

**Authors:** Thomas Folliard, Jerome Bonnet, John Ward, Frank Baganz, Chris Grant, Drew Endy

## Abstract

Boolean Integrase Logic gates use phage integrases that respond to transcriptional signals to implement logical functions within living cells via DNA recombination. Control of biological systems using these logic gates has many applications, including biomanufacturing, healthcare, or environmental remediation. Gates are highly sensitive to background transcriptional noise producing unwanted integrase expression and uncontrolled, permanent gate switching. Consequently, connecting gates to novel control signals requires time-consuming directed evolution of regions regulating integrase expression. Here we present an approach in which the activity of an alkane biosensor is tuned to match levels of existing integrase control signals. By adjusting AlkS expression, we tuned the transcriptional output of *P_alkb_* to match the output of existing integrase controllers. We successfully connected *P_alkb_* to two integrases with different transcriptional control requirements and demonstrated the original logic function was conserved. Our method complements directed evolution approaches to connect Boolean integrate logic gates to novel transcriptional sources and will ultimately facilitate the systematic *in silico* design of gates responding to various control signals.

## 1 Introduction

Reliable genetic Boolean Integrase Logic (BIL) gates were built using phage integrases responding to transcriptional control signals and producing the full range of logic functions (AND, NAND OR, NOR, XOR, XNOR) via DNA recombination (1–3). The BIL architecture relies on two modules: a logic element containing directional DNA regulatory sequences such as promoters and terminators, nested within integrase recognition sites, and a recombination control module embedding two phage integrases under tight orthogonal transcriptional control. Phage integrase-mediated inversion or excision of directional DNA elements gives rise to the full range of logic functions.

For every novel research or engineering application, the logic gates has to be controlled by particular transcriptional signals. Given the enzymatic and unidirectional nature of the integrases controlling gate switching, these devices are extremely sensitive to non-specific activation due to background transcriptional noise(4). In the current iteration of BIL gates, integrase expression is controlled by tightly regulated promoters exhibiting very low basal levels of expression and strong activation when the inducer signal is received(4). In addition, integrase expression levels are down regulated at the translational and/or at the post-translational level. Therefore, directly connecting novel promoters of interest to existing integrase expression cassettes are likely to result in spontaneous gate activation due to promoter basal activity levels above gate switching thresholds. In this context, methods allowing the routine connection of BIL gates to novel control signals in a reliable fashion are crucial and currently not available.

While predictive combinatorial design has worked for biosynthesis optimisation (5) the current method to connect logic gates to novel transcriptional sources involves the directed evolution of a regulatory region like the Ribosome Binding Site (RBS), Ribozyme insulators, the start codon, or a C-terminal Ssra degradation tag, alone or in combination (3, 6–8). Libraries are generated by PCR using degenerate primers, and several hundred library variants are then screened to obtain constructs switching only in the presence of the desired inducer molecule. This approach, while effective, can be time and resource consuming.

Here we sought to explore an alternative method that we call "level matching” achieved by placing BIL gates under the control of novel transcriptional sources. Our approach consists of matching the transcriptional activity of the novel source promoter with that of promoters used to control current instances of the gates, while keeping the same downstream RBS-Integrase design, and conserving a dynamic range of promoter activity sufficient to cross the integrase expression threshold level required for gate switching. This approach is complimentary to library screening of RBS strengths and provides an expanded toolkit when integrating further novel transcriptional signals into BIL gates.

## 2. Materials and Methods

### 2.1. Strains and plasmids

Plasmids and strains used in this study are shown in Table 1.

**Table 1:** Summary of failure modes and engineering solutions taken to tune the alkane biosensor for compatibility with the BIL gates. Schematics depicting the engineering solution taken to resolve each failure mode.

| Strain or Plasmid | Description | Reference or Source |
| --- | --- | --- |
| *E. coli* |  |  |
| DH5 alpha | Expression and cloning host |  |
| BL21 | Expression host |  |
| Plasmids |  |  |
| pSB50C1 | Dual controller plasmid | 2 |
| pSB50C2 | pTET:GFP reporter | 2 |
| pSB50C3 | pBAD:GFP reporter | 2 |
| pSB4A4 | AND gate | 2 |
| pSB4A5 | XOR gate | 2 |
| pSB50C6 | Alkane inducible GFP reporter; normal *alkS* expression | This study |
| pSB50C7 | Alkane inducible GFP reporter; normal *alkS* expression, JBEI RBS | This study |
| pSB50C8 | Alkane inducible GFP reporter; Low *alkS* expression | This study |
| pSB50C9 | Alkane inducible GFP reporter; Rhamnose induced *alkS* expression | This study |
| pSB50C10 | Single controller; Alkane induced Bxb1 integrase expression; normal *alkS* expression | This study |
| pSB50C11 | Single controller; Alkane induced Bxb1 integrase expression; low *alkS* expression | This study |
| pSB50C12 | Single controller; Alkane induced TP901-1 integrase expression; low *alkS* expression | This study |
| pSB50C13 | Single controller; Alkane induced Bxb1 integrase expression; Rhamnose induced *alkS* expression | This study |
| pSB50C14 | Single controller; Alkane induced TP901 integrase expression; Rhamnose induced *alkS* expression | This study |
| pSB50C15 | Single controller; Alkane induced TP901-1 integrase expression; low *alkS* expression; GTG start | This study |
| pSB4A16 | Integrase assay plasmid | This study |

### 2.2. Experimental procedures and data collection

#### Cloning and DNA assembly

Newly assembled plasmids were transformed by heat shock into NEB D5 chemically competent *E. coli.* A one hour recovery in SOC media preceded plating onto LB agar plates with the appropriate antibiotics. For re-transformation of plasmids obtained from a plasmid preparation of *E. coli* transformants, a 5 minute protocol was used (9). All assembly was conducted using Gibson isothermal assembly (10, 11). The following antibiotic concentrations were used: Ampicillin (50 μg/ml), Chloramphenicol (34 μg/ml), Spectinomycin (50 μg/ml).

#### Backbone

The main backbone for the study was pSB50C1 containing Bxb1 integrase under the control of P_TET_ and TP901 integrase under the control of P_BAD_, (ColE1 origin of replication, 50–70 copies, chloramphenicol resistance).

#### Native alks, PalkB modulated expression of bxb1

To generate pSB50C10, the inducible cassette containing the *P_alkb_* promoter and *alks* under control of the native promoter and RBS was amplified from pSB50C7.

#### Low alks modulated expression of bxb1

To generate pSB50C11, the inducible cassette containing the *P_alkb_* promoter and *alks* under control of the P_4_ promoter and BCD22 was amplified from pSB50C8.

#### Low alkS modulated expression of TP901

To generate pSB50C12, the inducible cassette containing the *P_alkb_* promoter and *alks* under control of the P_4_ promoter and BCD22 was amplified from pSB50C8.

#### Rhamnose modulated expression of alks for Bxb1 expression

To generate pSB50C13, the new inducible cassette containing the *alks* transcription factor under the control of the rhamnose promoter, BCD8 and the *P_alkb_* promoter were synthesised by Geneart.

#### Rhamnose modulated expression of alks for TP901 expression

To generate pSB50C14, the new inducible cassette containing the AlkS transcription factor under the control of the rhamnose promoter, BCD8 and the *P_alkb_* promoter amplified from pSB50C13.

#### Integrase assay plasmid

To generate pSB50C16, the inverted fluorescent genes and then Superfolder GFP-along with their promoters were synthesized by Geneart as a single construct. The backbone was amplified from pSB50C4 by PCR to linearize the plasmid between the biobrick prefix and suffix removing the AND logic element. The insert was then assembled into the backbone and sequence verified.

### 2.2 Growth conditions and assays

#### Standard seed culture growth conditions

Ten ml overnight cultures were inoculated from −80 °C stocks and grown in Nutrient Broth medium (Beef extract 3.0 g/L, Peptone 5.0 g/L) for 15 hours at 37 ^o^C, 250 RPM shaking, 25 mm shaking amplitude and appropriate antibiotics in falcon tubes.

#### Standard induction conditions

Overnight seed cultures were spun down, washed and re-suspended in 1% (w/v) tryptone, 1% (w/v) sodium chloride and 0.5% (w/v) yeast extract. Cultures were diluted to a concentration of 1/100 and grown for 8 hours under standard conditions (37^o^C, 250 RPM) in the presence of an inducer. The following concentrations of inducer were used: anhydrous tetracycline 500 ng/ml, L-arabinose 0.1% W/V, DCPK 0.05% V/V.

#### Growth measurements

Overnight triplicate cultures were diluted to a 1/1000 concentration and placed with relevant antibiotics in Greiner 96 deep well plate plates.

#### Measurement of fluorescence intensities

Expression of the Superfolder GFP (14) or mKate2 (15) was measured using a GENios plate reader (Tecan) at an excitation wavelength of 485 nm and an emission filter of 520 nm for SuperfolderGFP and an excitation wavelength of 580 nm and an emission filter of 630 nm for Mkate2 using a Greiner black 96 well plate. Fluorescence was normalized by optical density using the following equation:

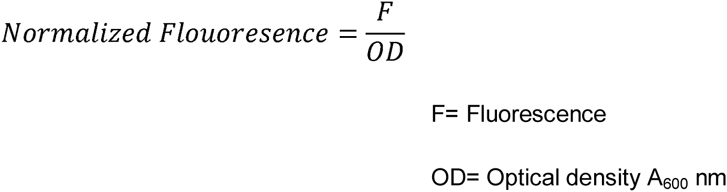

Flow cytometry measurements were obtained using a Beckman Coulter Epics XL Flow Cytometer. The results were analysed using the Summit Flow Cytometry analysis software (Beckman Coulter).

#### Integrase assay

Integrase expression was measured using the integrase assay plasmid (pSB4A16). The construct containing the integrase under control of a test cassette was co-transformed into chemically competent DH5*α*Z1 *E. coli* cells(12) along with the pSB4A16 plasmid. Cells were then grown in LB at 37 ^o^C in 0.5 ml cultures in Greiner deep, 96 well plates for 8 hours in the presence of stated inducers and shaken at 1000 RPM. The full 0.5 ml volume was taken from each test sample, spun down at 13,000 RPM and washed in PBS.

## 3. Results and discussion

As a model system, we used an alkane-sensitive promoter derived from the OCT plasmid in *Pseudomonas putida* GPo1 (13–15). In its native context, the AlkS system triggers expression of an alkane degradation pathway allowing for utilisation of alkanes as a carbon source for growth (16–18).

To place the logic gates under the control of the alkane biosensor, we measured the P_TET_ and P_BAD_ promoter transfer functions via a GFP assay and used these curves as a reference standard to which *P_alkb_* expression was then modulated. We reduced the basal activity of the *P_alkb_* promoter until its output was indistinguishable from that of the reference promoters. The resulting biosensor system was then successfully connected to the logic gates. We found that reducing the complexity of the ALKS feedback system and using standard biological parts from the BIOFAB library (19) significantly facilitated the engineering process. Our method complements directed evolution approaches to connect Boolean integrate logic gates to novel transcriptional sources.

### 3.1 Design and characterisation of a novel Integrase activity reporter.

Previously published logic gates did not support independent measurement of integrase activity (1, 2, 4). Thus, to facilitate our assays, we first devised an approach where controlled switching for both integrases was independently measured using the same construct (FIG 1A). We nested genes encoding two fluorescent proteins with distinct excitation-emission spectra, a Green Fluorescent Protein, Superfolder GFP (20), and a Red Fluorescent Protein, mKate2 (21), in inverted orientations within integrase recognition sites. Inversion of the fluorescent protein sequence allowed for translation of the forward, correct open reading frame and for successful expression. Using this plasmid, we were able to rapidly and independently assess the function of the two integrases Bxb1 and TP901 in the presence of their cognate inducer (FIG 1B). Characterisation of these constructs provided a direct assay for Integrase mediated DNA inversion.

**Figure 1:**
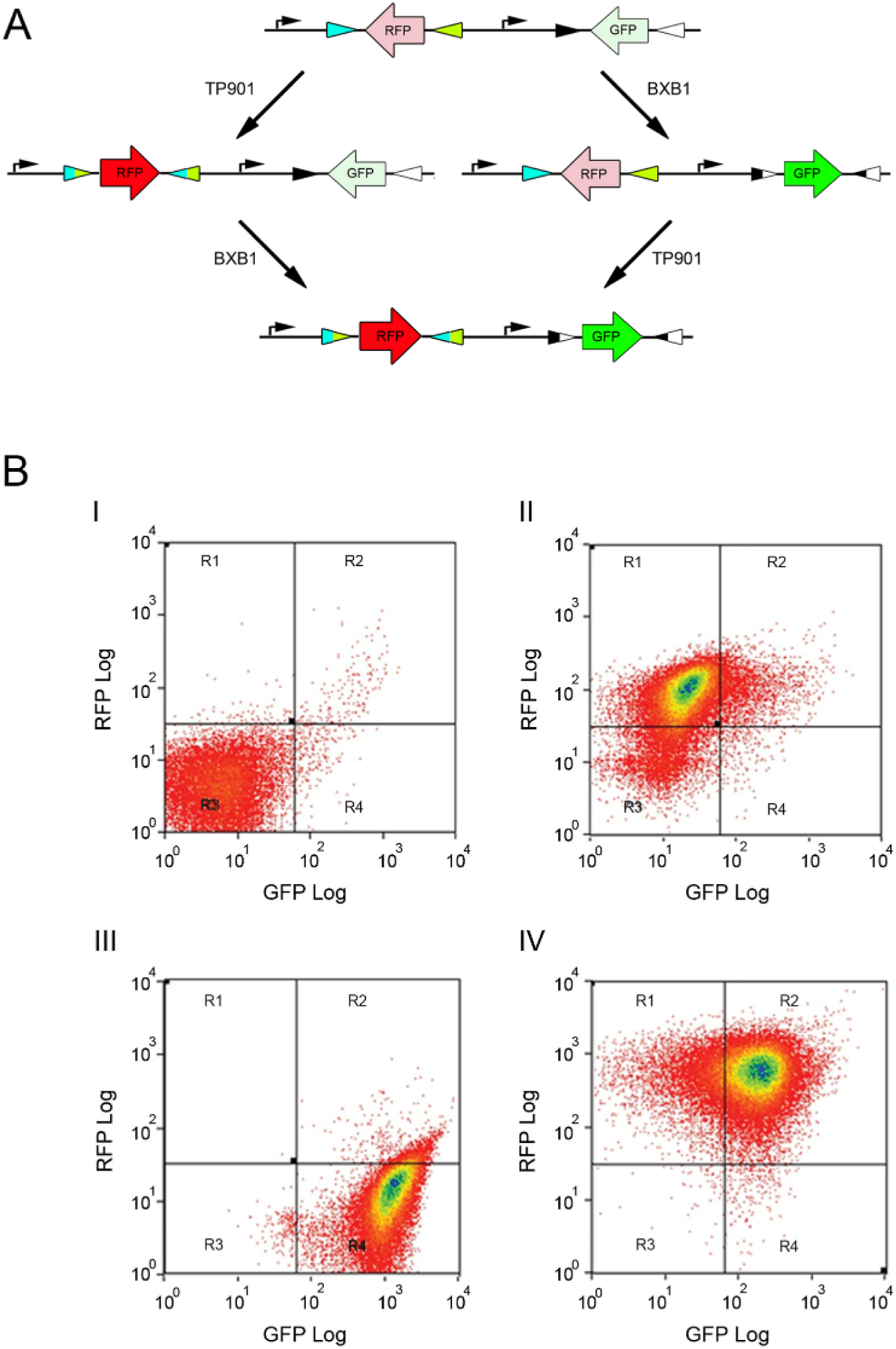
Design and characterisation of the integrase dual reporter plasmid. **(A)** Architecture and operational principle of the dual reporter plasmid for independent measurement of integrases activity. Coding sequences for two different fluorescent proteins (superfolder GFP and mkate2) are nested between recombination sites from independent serine integrase (Bxb1 and TP901–1, respectively). Both fluorescent proteins are placed in reverse orientation compared to the transcriptional flow coming from a constitutive promoter, leading to the transcription of a non-coding RNA and no fluorescence in the absence of integrase mediated switching. The TP901 integrase mediates inversion of the mkate2 coding sequences, allowing for RFP expression. Bxb1 mediated inversion of the Superfolder GFP coding sequence leads to production of GFP. When both integrase are active, both fluorescent reporters are produced. **(B)** Experimental characterisation of the dual reporter plasmid by flow cytometry. (i) no inducer. (ii) Anhydrous tetracycline controlling Bxb1 expression (500 ng/ml). (iii) L-arabinose 0.1% w/v controlling expression of TP901 and (iv) Both Anhydrous tetracycline 500 ng/ml and L-arabinose 0.1% w/v.

### 3.2 Development of an alkane responsive biosensor.

We then characterised the transfer function of the alkane biosensor in response to dicyclopropylketone (DCPK), an alkane mimic (22, 23). We amplified the full AlkS coding region and the *P_alkb_* 5′ UTR. We replaced the first open reading frame in the alkane degradation pathway *–alkb-* with the gene encoding for GFP (FIG 2A and Fig S2A).This kept expression of GFP under the control of the wild-type *alkb* RBS. However, we found that this construct failed to respond to DCPK (FIG 2B), analysis of the full *P_alkb_* 5′ UTR along with the first 100 base pairs of the GFP open reading frame through the RNAFold software (24) using a minimal free energy model (25) showed the lack of expression was due to *de novo* secondary structure introduced between the wild type RBS (ΔG:-19.42) and the new ORF (ΔG:-18.92) (S2 B). We introduced a new RBS with no predicted secondary structure and then experimentally confirmed that the new construct, the *P_alkb_*:GFP reporter, produced the expected induction response after DCPK addition (FIG 2C). These data demonstrated a responsive, *in vivo,* alkane biosensor prior to introduction into the BIL gates.

**Figure 2:**
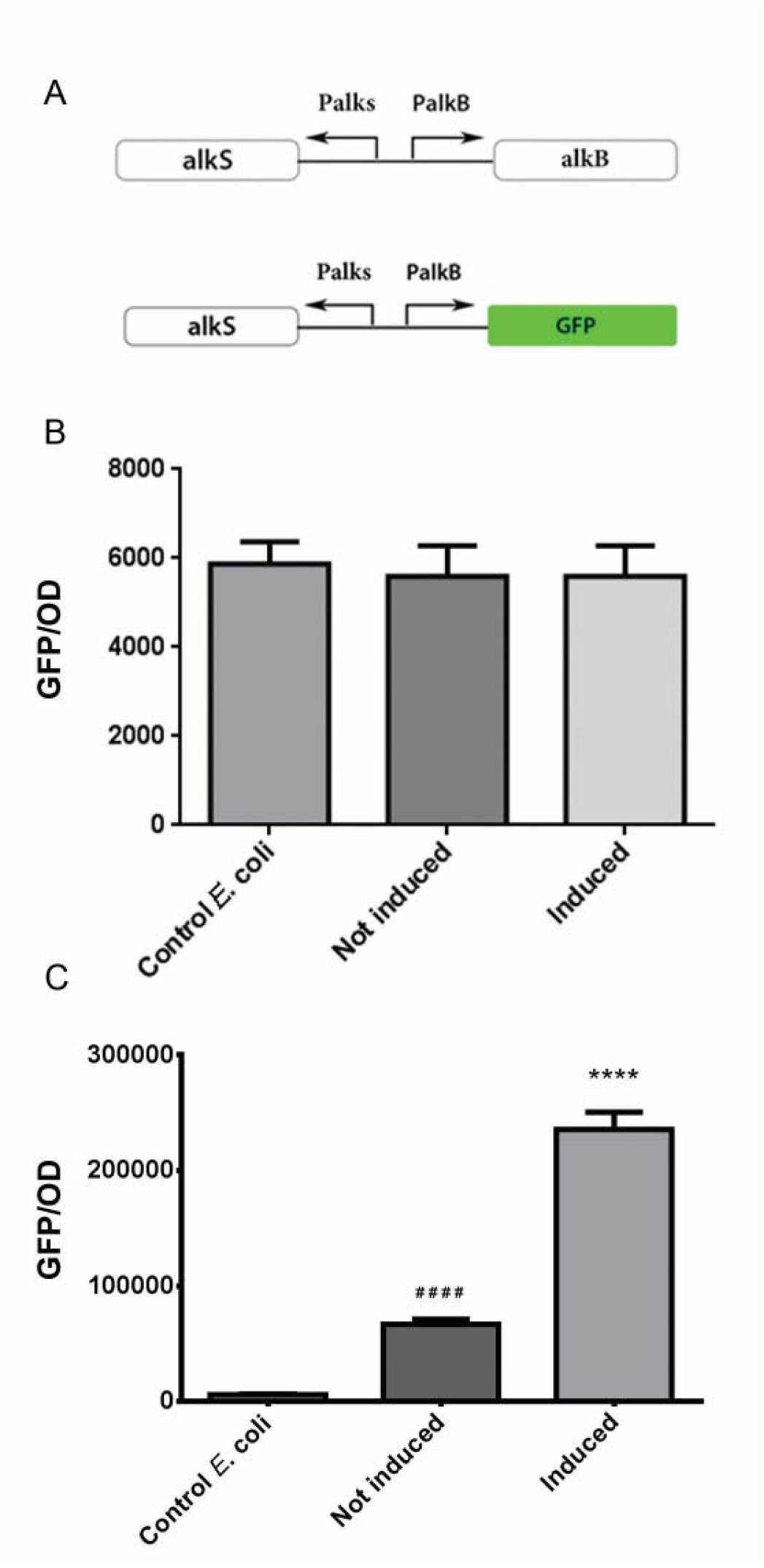
Construction and characterisation of an alkane biosensor derived from the *Pseudomonas putida* ALK operon. ***(A)*** Schematics of the Alkane biosensor reporter. The first enzyme in the alkane degradation pathway (*AlkB* gene), was replaced by GFP. The native *AlkB* RBS was conserved ***(B)*** Lack of response to DCPK for a biosensor in which the native RBS controls GFP translation. No statistically significant difference was observed between negative control *E. coli* cells and the construct either in the presence or absence of the DCPK inducer (0.05% v/v, triplicate repeats). ***(C)*** Induction response from the alkane biosensor with a newly designed RBS. Addition of DCPK (0.05% v/v) produced a 3.5 fold increase in iGFP intensity compared to non-induced cells. Fluorescence level in non-induced cells due basal expression of GFP from the alkane biosensor was 11 times higher than in the control *E.coli* strain. For all panel, (****) indicates paired t-test against control *E. coli*, P<0.0001 (Experiments were performed in triplicate and error bars indicate standard deviation).

### 3.3. High basal activity of the alkane biosensor leads to spontaneous Bxb1 mediated gate switching.

We then replaced the Bxb1 integrase control signal, P_TET_, by the alkane biosensor. In our design, we kept Bxb1 original RBS (FIG 3 A). We found that basal Bxb1 expression from the alkane biosensor was high enough to produce spontaneous flipping in the absence of any inducer (FIG 3 B). Using the P*_alkb_*:GFP reporter, we found that the alkane biosensor had a basal expression level ~ 6 fold higher than the P_TET_ promoter (FIG 3 C). The main cause of high basal expression was compounded due to a gene dosing effect caused by the overexpression of the ALKS transcription factor from a high copy plasmid, rather than the very low copy number OCT plasmid from which the AlkS system originates. These data demonstrate that differences in expression levels (or level mismatch) is a main challenge to be overcome in order to directly connect novel transcriptional sources to Boolean Integrase Logic gates.

**Figure 3:**
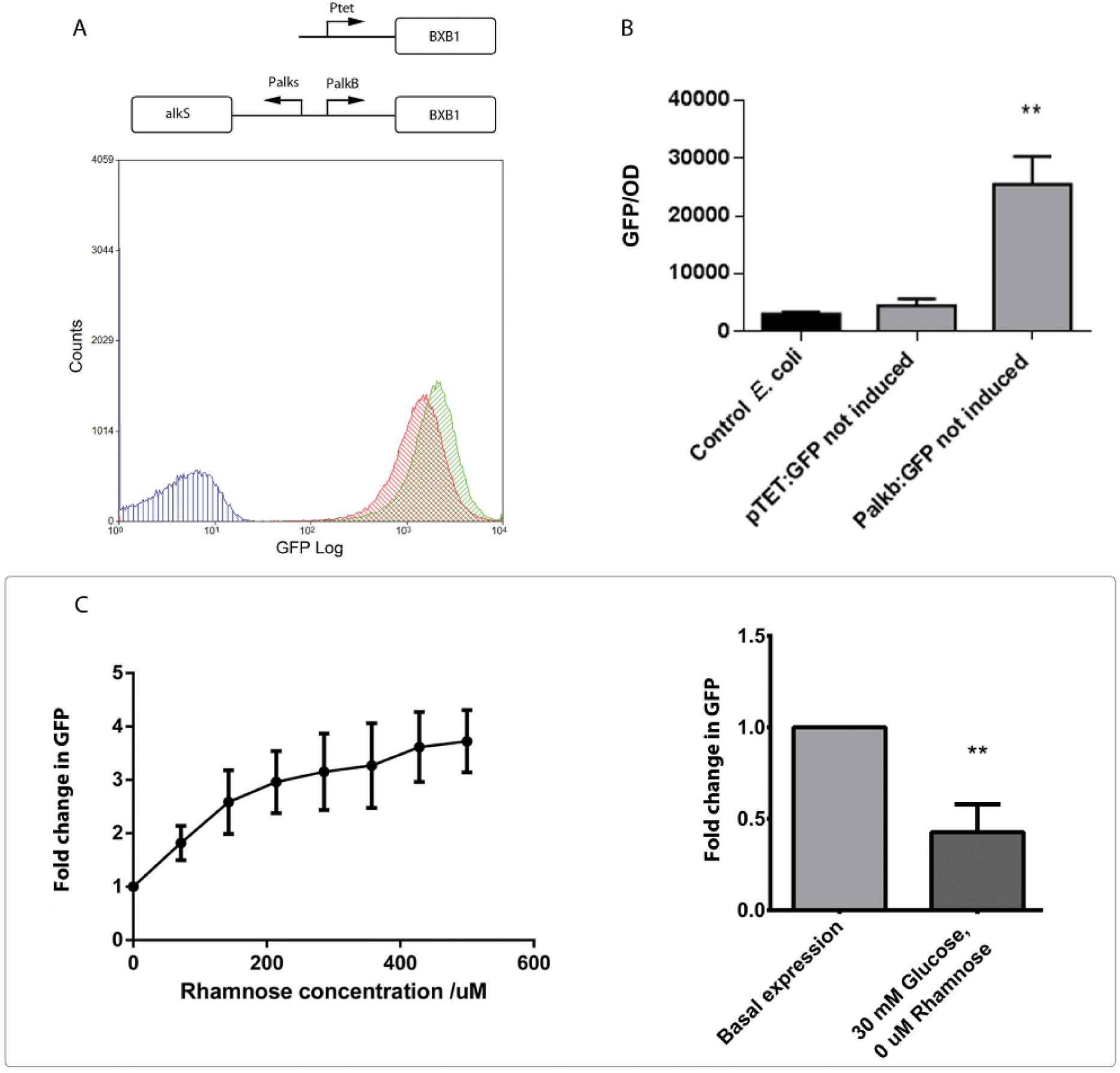
Elevated basal expression from the alkane biosensor leads to Bxb1 mediated DNA inversion in the absence of inducer. ***(A)*** Schematics of the constructs in which the alkane biosensor drives Bxb1 expression. The *P_TET_* promoter was replaced by the alkane biosensor while Bxb1 remained under control of its original RBS. Spontaneous Bxb1 mediated DNA inversion in the absence of DCPK. Flow cytometry analysis of induced and uninduced colonies. Induced cells (red histogram) -DCPK 0.05% v/v; uninduced cells (green histogram), and control cells (blue histogram)-no plasmid. No discernible difference between induced and uninduced cultures indicates that basal Bxb1 expression is sufficient to promote flipping. ***(B)*** Bulk population comparison of basal GFP expression from the *P_TET_* and the *P_alkb_* promoters. Basal expression from the *P_alkb_* promoter was 6 fold higher than *P_TET_* promoter. ***(C)*** Effect of modulating ALKs expression on induction and basal expression from P*_alkb_*. *Left panel:* **c**hange in response to 0.05 V/V DCPK with rising rhamnose concentration measured using the P*_alkb_*_-_GFP reporter. For every data point, fold change is calculated relative to basal expression level. *Right panel:* effect of glucose repression on *P_alkb_* activity expression showing a two fold reduction in basal expression from the *P_alkb_* promoter (Experiments were performed in triplicate and error bars indicate standard deviation).

### 3.4. Matching *P_alkb_* and *P_TET_* basal activity level.

We therefore needed to reduce P*_alkb_* basal expression by decreasing the expression of the P*_alkb_* transcription activator, ALKS. To find the suitable ALKS expression levels, we replaced the native ALKS promoter with the inducible rhamnose promoter. We chose to use the rhamnose promoter for two reasons: (i) a rheostatic control of gene expression levels allowing us to modulate the intracellular levels of ALKS and putatively the transcription activity of *P_alkb_* and (ii) the possibility of epressing expression through glucose catabolite repression due to a cAMP binding site positioned upstream of the promoter site (if the basal expression of the rhamnose promoter would still be to high). To find a suitable RBS, we used the BIOFAB collection (26), a library of standardised transcriptional and translational control elements which offers a ladder of expression levels to choose from, independently of the downstream gene sequence. We selected the BCD8 RBS which provides a wide translation range when transcription levels are modulated.

Using a P*_alkb_*-GFP reporter, we confirmed that varying concentrations of rhamnose could modulate P*_alkb_* transcription output: increasing rhamnose concentrations allowed for a greater response to the same levels of DCPK inducer. Moreover, we found that addition of 30mM L-Glucose gave a statistically significant reduction in basal GFP expression (Fig 3 D). Based on these results, we concluded that we needed to place the ALKS regulator under a weak transcriptional control. From the BIOFAB collection we identified a weak constitutive promoter and a weak bi-cistronic RBS and replaced the *alks* 5′ UTR with the P4:BCD22 pair (BCD22 being one of the weaker bi-cistronic RBS of the library). The P4:BCD22-ALKS construct produced a basal expression statistically indistinguishable from the original *P_TET_* architecture (FIG 4 A).

**Figure 4:**
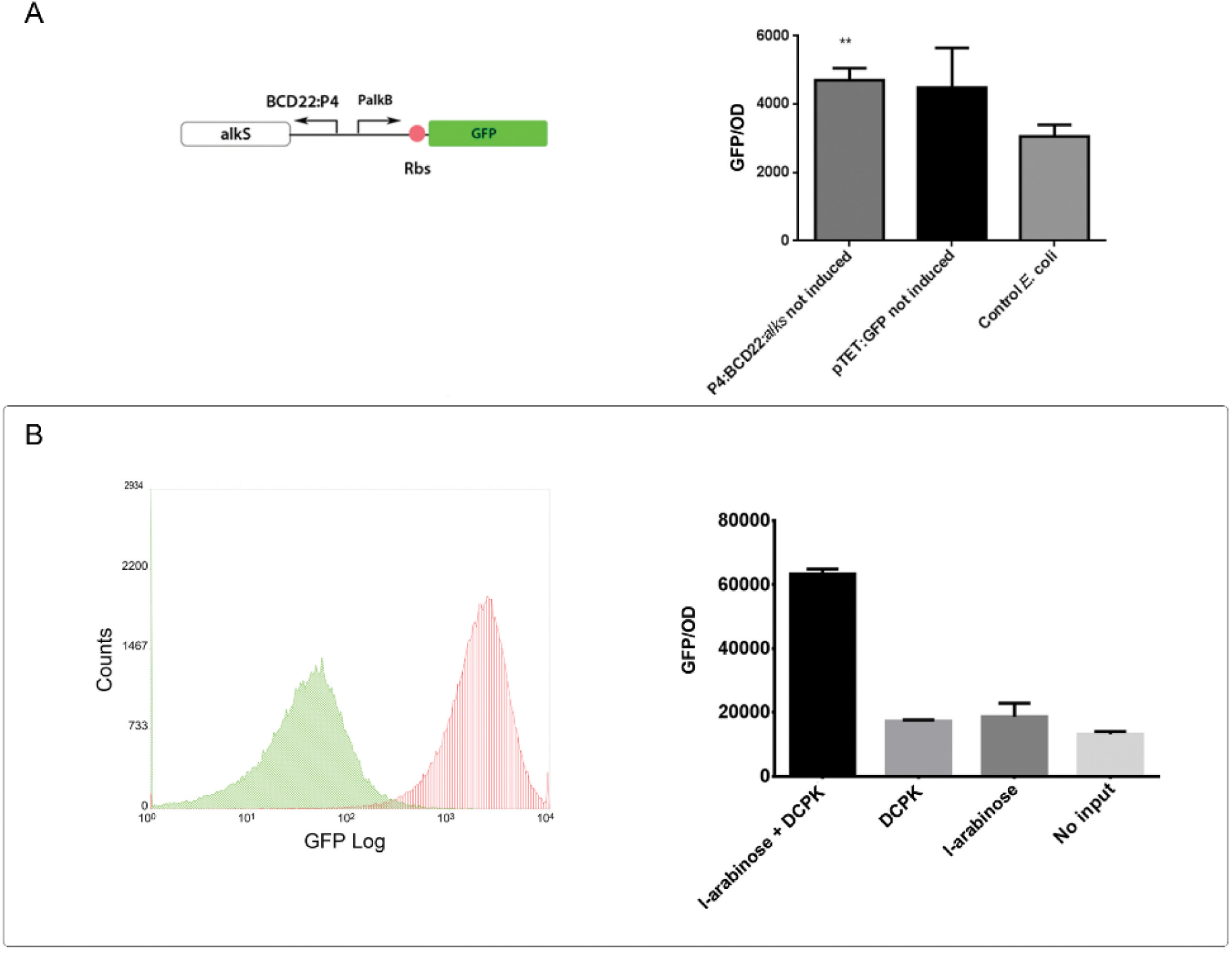
Using the level matching approach to place Bxb1 under the control of the alkane biosensor. **(A)** Modulation of the alkane biosensor transcriptional output. Replacing the dual promoter architecture from the AlkS transcription factor with the weak P_4_-BCD22 promoter/RBS pair from the BIOFAB library reduce basal expression of the alkane biosensor close to the levels of the pTET promoter. **(B)** Alkane induced expression of BXB1. *Left panel:* Flow cytometry measurement of cells containing the Integrase assay plasmid showing a distinct induction response in the presence of 0.05 V/V DCPK (red histogram) compared with uninduced cells (green histogram). *Right panel:* Response of the AND logic gates - on plasmid pSB4A4-showing both DCPK and L-Arabinose are required to produce GFP. For all panels: ** represents Paired T-test P values P<0.01, (Experiments were performed in triplicate and error bars indicate standard deviation).

We then placed the Bxb1 integrase under control of the P_4_:BCD22 modulated biosensor and found that no spontaneous switching was observed in the absence of DCPK. Moreover, induction with DCPK triggered a transcriptional response and enabled the AND gates to function as predicted (FIG 4 B). We also tested whether the use of a bi-cistron might confer some toxic effects from expression of the leader peptide or nonsense protein. Growth measurements of cultures containing the unmodulated and BCD8/22 modulated biosensors showing no toxicity (Fig S3). Taken together, these results demonstrate that Boolean Integrase Logic gates can be directly placed under the control of novel transcription sources whose basal expression level has been matched with the basal expression of known gates control signals. These data demonstrate that modulation of the ALKS transcriptional activator allowed for modulation of the *P_alkb_* promoter in such a way that its activation levels became indistinguishable from the original P_tet_ promoter used in the BIL gates to control BXB1 integrase and that this was sufficient to successfully integrate the novel transcriptional source into the BIL gates.

### 3.5 Using the level matching approach to replace TP901 control signal with the alkane responsive biosensor.

As replacing the *alks* 5’ UTR with P_4:_ BCD22 combination allowed for successful introduction of Bxb1 under alkane biosensor control, we then sought to validate our method using another integrase as different enzymes have different activities and stabilities and therefore, depending on the enzyme, gate switching will occur at different transcriptional threshold. Additionally, the basal transcriptional level causing spontaneous, non-induced switching will vary.

We thus placed the TP901 integrase under the control of the *P_alkb_* promoter regulated by the P_4_:BCD22-ALKS cassette. As with Bxb1, we conserved the TP901 cassette, including the RBS and a C-terminal SsrA degradation tag (5). However, unlike with Bxb1, we failed to successfully modulate TP901 expression and spontaneous flipping occurred in the absence of any inducer (Fig. S4 A,B), indicating that the TP901 integrase cassette had different transcriptional control requirements than the Bxb1 cassette. Further analysis revealed that the P_BAD_ promoter originally controlling TP901 had a statistically significant lower basal expression than the BCD22 modulated biosensor, indicating that further reduction of basal integrase expression was required (S4 C).

We changed the TP901 start codon from an ATG to a GTG reducing its expression to a third (18) and observed a reduced basal expression sufficient to prevent spontaneous DNA inversion. However, there was no further response to the inducer indicating that expression had been reduced too strongly. To identify an *alkS* expression level for which the basal transcriptional activity is below the threshold needed to activate TP901 yet sufficient to enable flipping in the presence of the alkane/DCPK control signal, we again used the rhamnose controlled ALKS expression vector in combination with the connected P*_alkb_* promoter controlling to the original TP901 cassette (FIG 5 A). We found that basal expression of ALKS from the rhamnose promoter in the presence of glucose repression was sufficient to prevent TP901 spontaneous switching while allowing for reliable TP901 mediated inversion in the presence of 0.05% DCPK inducer (FIG 5 B). Therefore, we successfully connected TP901 to a novel transcription source. These data validate the generic nature of our level matching approach by applying it to a different integrase cassette having different transcriptional control requirements.

**Figure 5:**
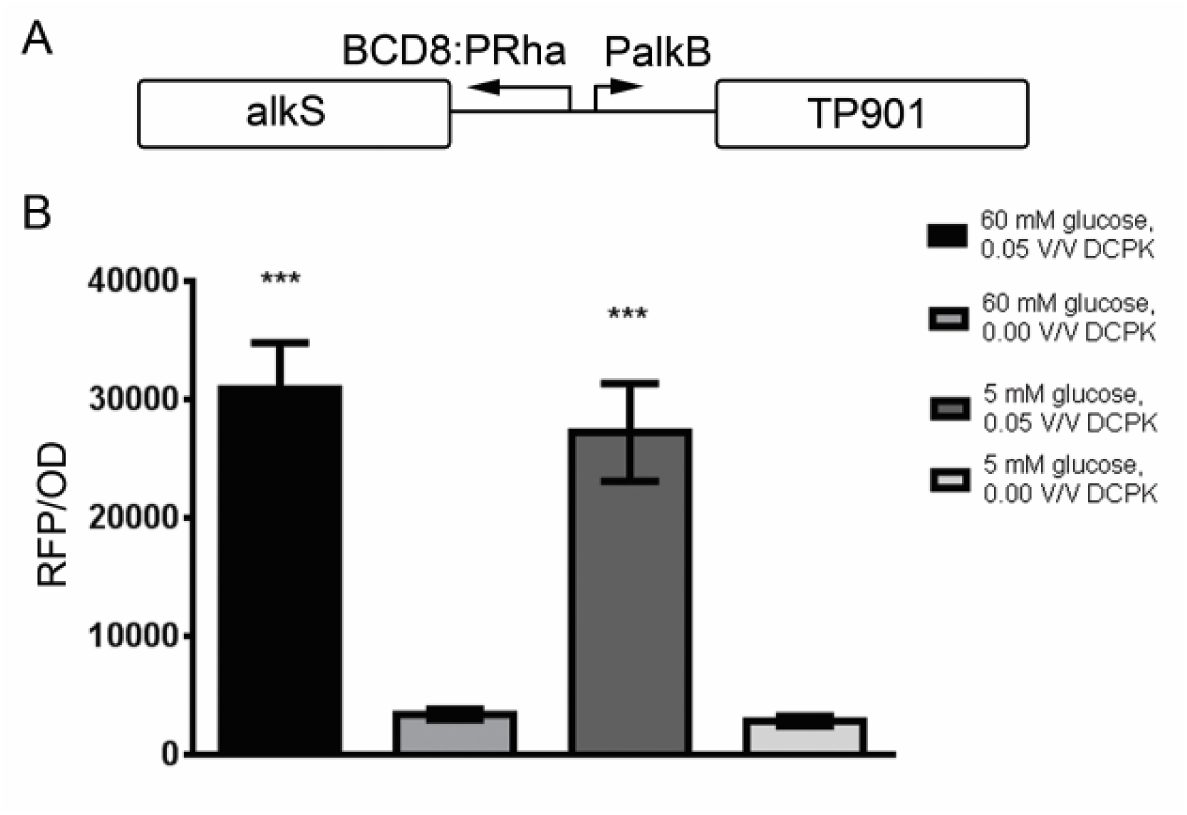
Using the level matching approach to place TP901 under the control of the alkane biosensor. **(A)** Re-modulation of the alkane biosensor using the rhamnose promoter and a wide range RBS–BCD8. **(B)** Induction response to 0.05% V/V DCPK showing reliable TP901 mediated flipping of RFP in the presence of 60 and 5 mM glucose (*** indicates paired t-test against not induced culture, P<0.001, experiments were performed in triplicate and error bars indicate standard deviation).

## 4. Conclusion

Here we demonstrate a novel methodology to introduce a biosensor of interest as a control signal for Boolean Integrase Logic gates. We successfully replaced both P_TET_ and P_BAD_ control signals by the P*_alkb_* alkane biosensor and reliably activated both integrases (TP901 and Bxb1) in response to the new input signal while avoiding spontaneous switching. To do so, we modulated the activity of the ALKS transcription factor to match the transcriptional control levels for two different integrases requiring different levels of basal expression and activation to operate.

We first defined the Bxb1 and TP901 input thresholds by measuring the ALKS activator expression level required to avoid spontaneous switching while still supporting switching in the presence of inducer. Using the inducible Rhamnose promoter to control the levels of the transcription activator ALKS controlling integrases expression allowed us to map the transcriptional space of the P*_alkb_* promoter activities more efficiently. We then used the BIOFAB dataset to choose the appropriate combination of promoter and BCD. Our work thus also highlight the importance of standardised, well-characterised and open source libraries of biological parts.

Promoter strength matching by comparing a desired transcriptional source output with the output of the P_TET_ or P_BAD_ promoters controlling integrase expression and then adjusting the output of this novel transcriptional source to be statistically indistinguishable from the original promoters could thus be used as a generic approach to introducing novel control inputs into the BIL gates without the need for directed evolution. A summary of the engineering solutions taken to tune the alkane biosensor for compatibility with the BIL gates is shown in Table 1. An overview of the approach taken to introduce the alkane biosensor into the BIL gates is shown in S4.

Our work presents a rational alternative to small scale screening of RBSs or directed evolution approaches and removes the need for library generation and screening efforts and could hence take considerably less time and cost.

Once a specific gate control signal range has been determined and given the availability of well characterised libraries of controllers offering a range of different strengths, our method should be amenable to the *in silico* design of gates with novel control signal in a more predictable way. Such libraries of standardised and well-characterised regulatory elements are already available and their number should be increasing in the next few years, allowing for a rapid, computer-assisted level-matching approach to generating logic gates responding to novel control signals, facilitating the deployment of Boolean Integrase Logic for many applications.

## Acknowledgements

We wish to thank J, Armitage, T. Bayer, T. Call, S. Damalas, M. Gershater, J. Hales, C. Harrison, J. Jefferies, B. Lichman, L. Malivert, B. Mertins, R. Rodrigues, L. Rothschild, J. Rutley, V. Sanchania, R. Young and members of the Bayer, Endy and Ward group.

## SUPPLEMENTARY DATA

**S1.**
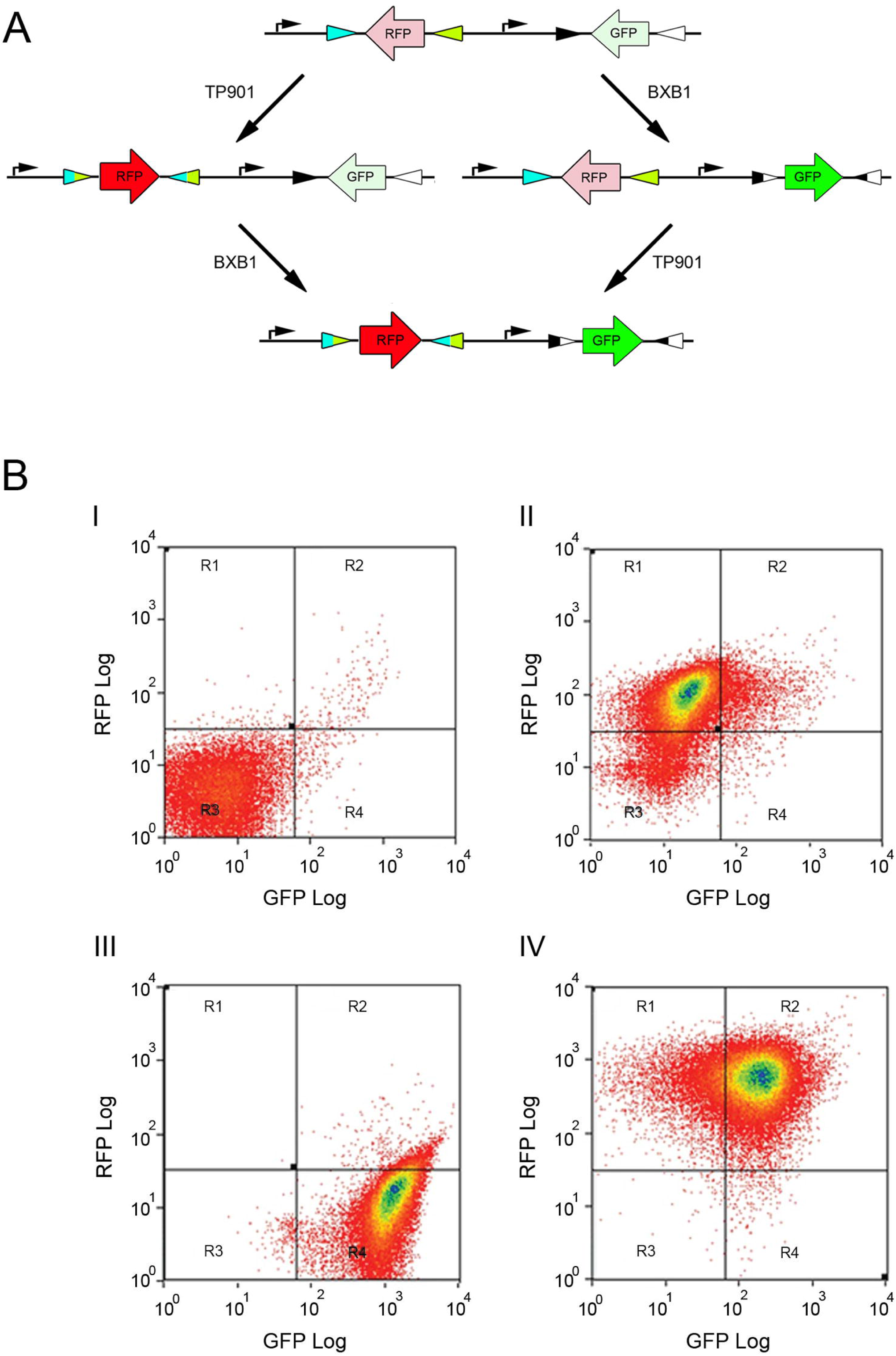
(A) Superfolder GFP replaced AlkB on an ATG for ATG basis keeping the 5' UTR unchanged with the intention of translating SFGFP from the wildtype RBS. (B) Predicted secondary structure around the RBS indicates the introduction of new secondary structure. Predicted secondary structure indicated by parenthesis. Introduction of a new RBS removed the predicted secondary structure between the RBS and SFGFP.

**S2.**
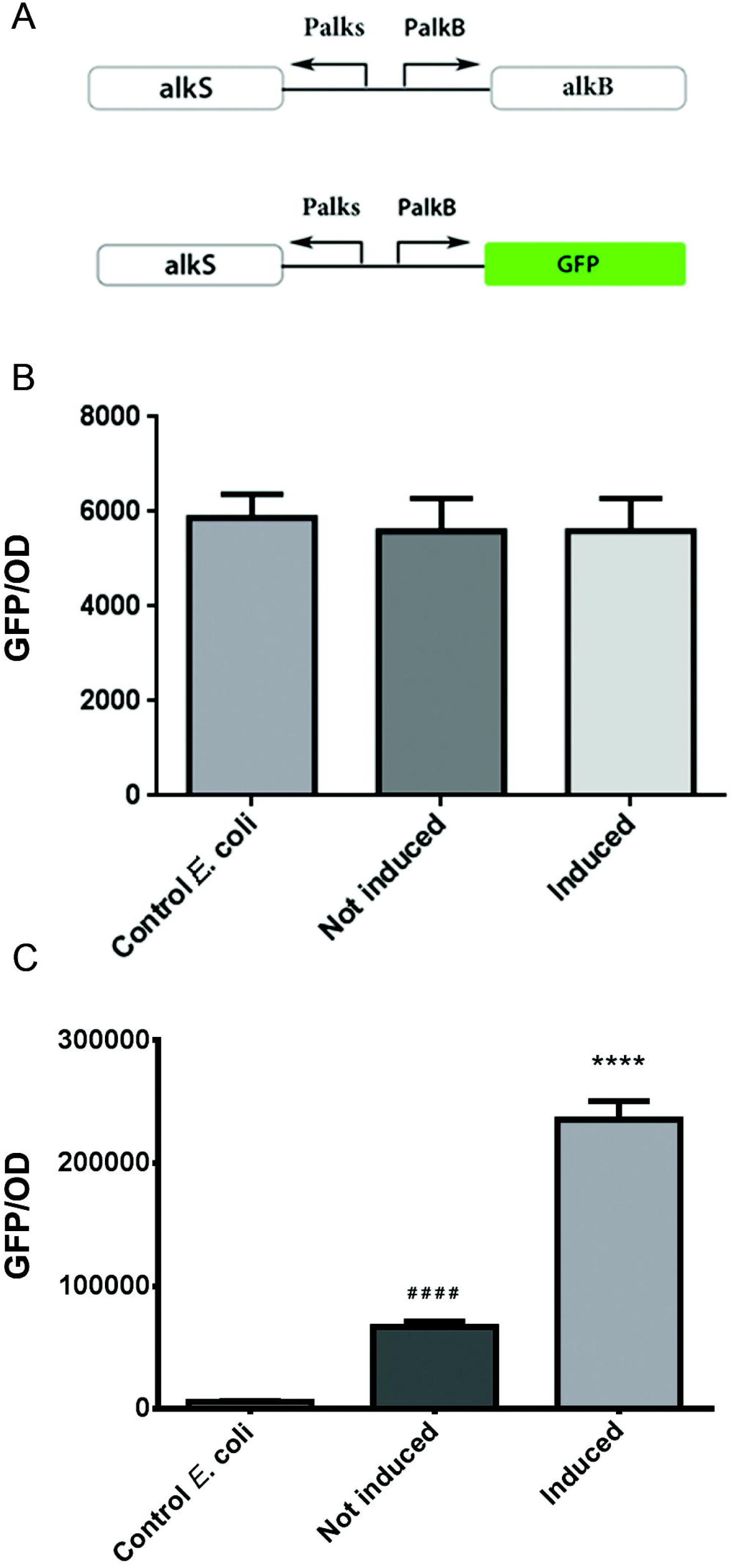
Effect of Bi-cistronnic RBS BCD22 on growth rate. A) growth rates from the BCD22:P_4_ modulated (pSB50C8) and unmodulated biosensor (pSB50C7) showing an increased growth rate most likely due to reduction in burden from alkS over expression. B) growth rates from the BCD22:P_4_ modulated (pSB50C8) and P_RHA_ (pSB50C9) showing no significant change in growth between bi-cistron 8 and 22. C) Overlay of all three constructs showing an improved growth rate conferred from the use of a bi-cistron indicating no toxic effect that was not compensated for by reduced burden on the cell.

**S3:**
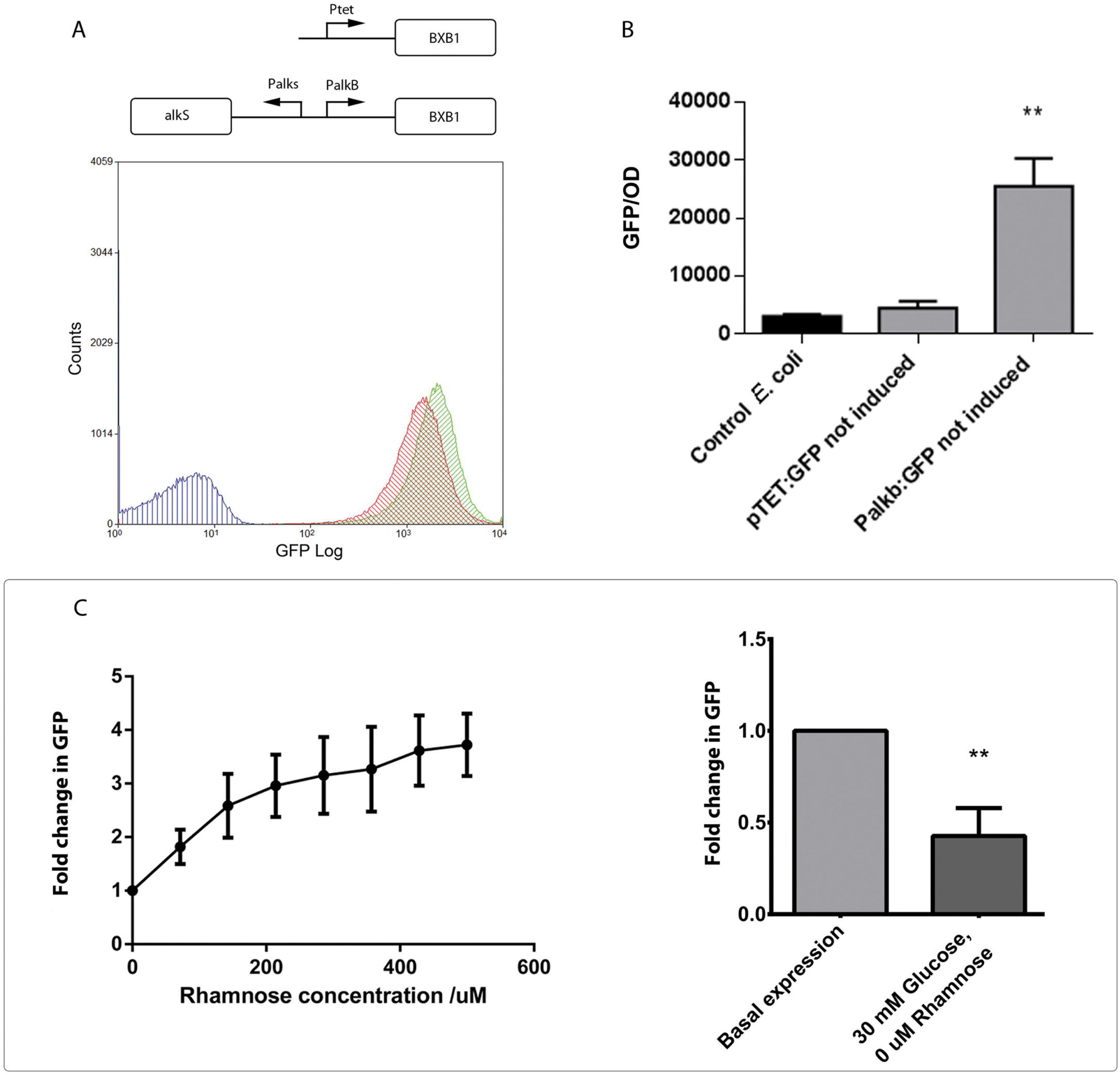
Introduction of TP901 Integrase under control of the P4:BCD22 modulated biosensor. A) TP901 Integrase under control of the P4:BCD22 modulated alkane biosensor. B) Basal expression from the assay plasmid - pSB4A16-showing a high basal RFP expression showing TP901 has been expressed sufficiently in the absence of any inducer to reliably invert its target.

**S4:**
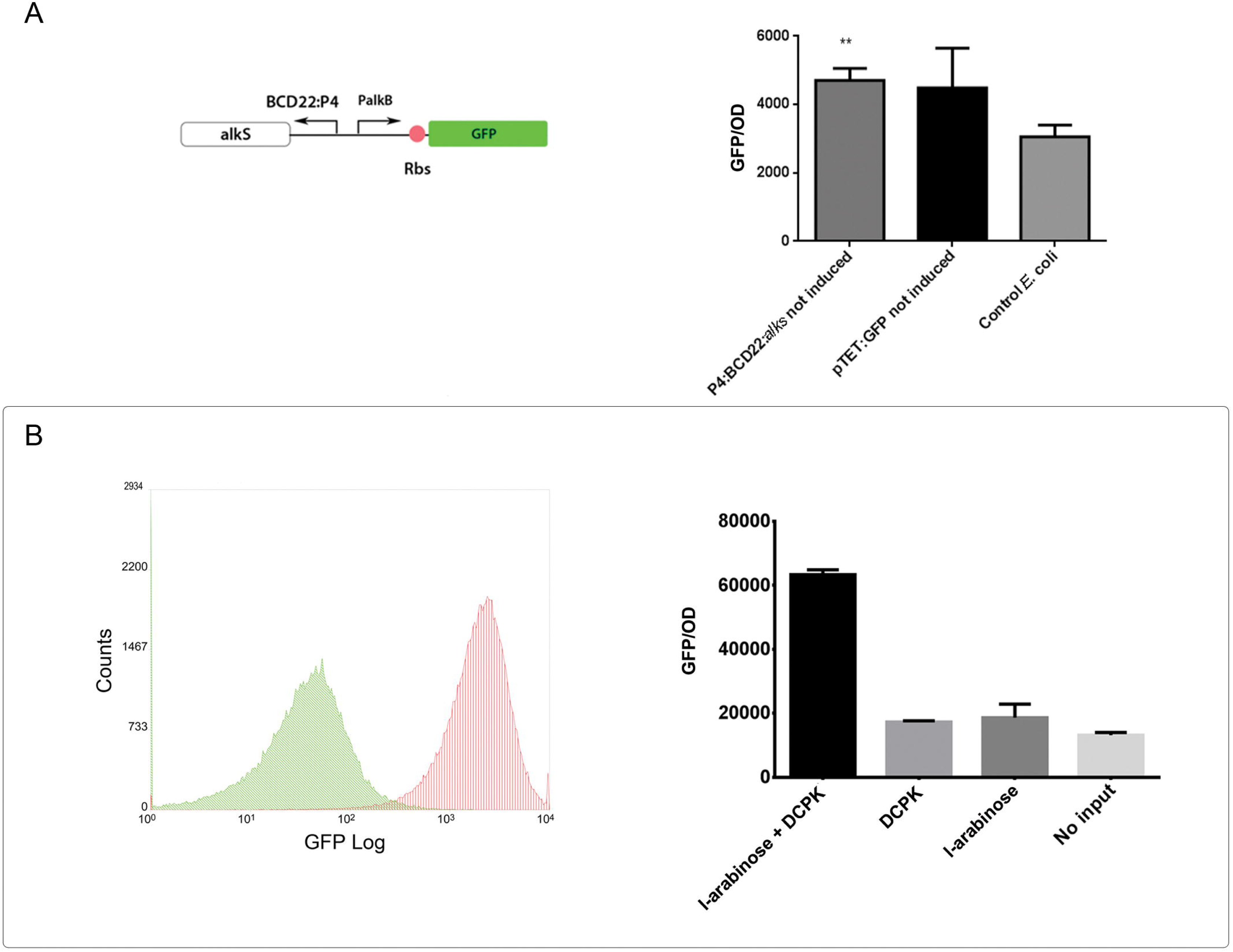
list of plasmids used in this study

**S5:**
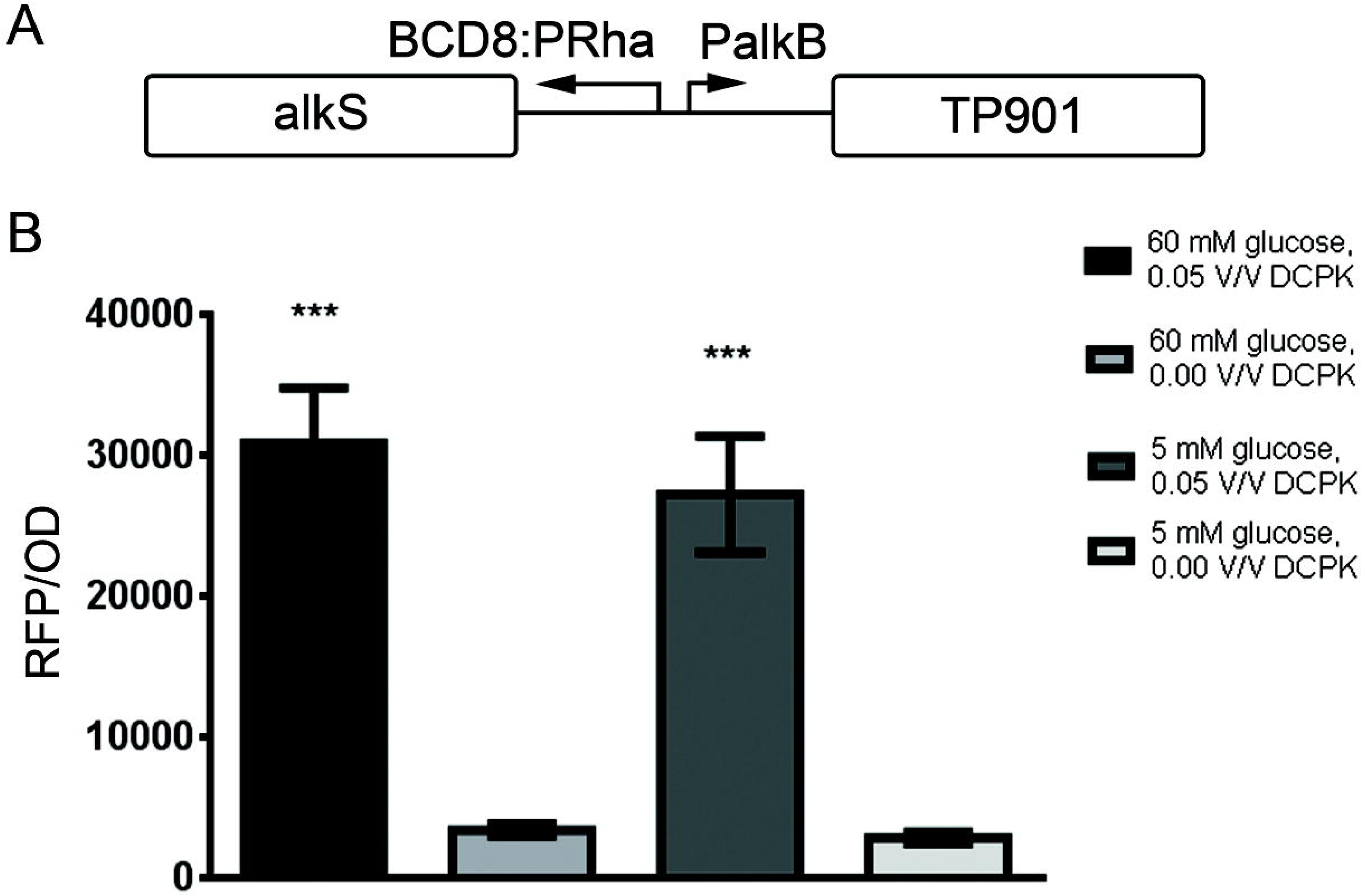
Overview of the approach taken to BIL gate re-engineering.

